# Predator recognition and anti-predatory behaviour in a recent aquatic invader, the killer shrimp (*Dikerogammarus villosus*)

**DOI:** 10.1101/636100

**Authors:** Matteo Rolla, Sonia Consuegra, Carlos Garcia de Leaniz

**Affiliations:** Department of BioSciences, Centre for Sustainable Aquatic Research, Swansea University, Singleton Park, Swansea SA2 8PP, UK

**Keywords:** Invasive species, Anti-predatory strategy, Chemical recognition, Aggregation behaviour, Killer shrimp

## Abstract

The killer shrimp (*Dikerogammarus villosus*) is one of the most recent, but also most damaging, aquatic invasive species in Europe, but information on how the species responds to novel predation pressures in recently invaded areas is very limited. We employed an open test arena to examine predator recognition and anti-predatory behaviour in killer shrimp exposed to either blank water or water conditioned with fish kairomones to simulate a predator threat. Within five years after their introduction, killer shrimp spent much more time hiding in the presence of fish kairomones than when they were exposed to blank water. However, no significant difference was found in aggregation behaviour, and killer shrimp were strongly attracted to the scent of conspecifics regardless of predator threat. Given the strong selective pressures that fish predators can exert on native and invasive gammarids, our findings highlight the need to consider prey-predator interactions to better predict the dispersal and likely impact of killer shrimp into invaded ecosystems.

## Introduction

From a prey-predator perspective two opposing selective forces may confront invasive species when they colonise a new area: the absence of former predators may facilitate their establishment (the enemy release hypotheses – Colautti et al. (2004), while their different appearance (the oddity effect - Almany et al. (2007) and lack of co-evolutionary history (the ‘naïve prey’ hypothesis - Sih et al. 2010) may curtail it. Thus, whether invasive species thrive or flounder may depend on what predators they encounter, and how they respond to them. This may result in ‘boom and bust’ cycles, reflecting prey-predator dynamics (Strayer et al. 2017). Surprisingly, very little is known about anti-predatory strategies of invasive species in novel habitats.

The killer shrimp (*Dikerogammarus villosus*) is a freshwater gammarid indigenous to the Ponto-Caspian region which has recently invaded Western Europe (Tricarico et al. 2010), and which therefore constitutes a good system to examine anti-predatory strategies in novel habitats. The species has a small size (1.8-30mm; Aldridge 2015) a flexible omnivorous diet (Mayer et al. 2008), and lives in a wide variety of freshwater and brackish habitats (Devin & Beisel 2008) where it faces many different potential predators. Despite its very recent introduction, it is listed among the 100 most invasive species in Europe (DAISIE 2009) and included in the RINSE (Reducing the Impact of Non-Native Species in Europe) black list with a score of 9 out of 10 (Gallardo et al. 2016). It can displace and prey on local gammarids and reduce native biodiversity (Eckmann et al. 2008; Macneil et al. 2013), and may already be benefitting from a boom phase in some parts of Europe, having shed some of its former parasites (Arundell et al. 2015; Grabner et al. 2015). The need for more information on this aquatic invader has been flagged as a priority (Gallardo et al. 2016; Pöckl 2009), as it is predicted that the species will cause major deleterious impacts on native fauna (MacNeil et al. 2012).

Studies on the killer shrimp have focused mostly on its diet, there is very little information on its predators. The species appears to be trophically very plastic (Bacela-Spychalska & van der Velde 2013; Casellato et al. 2007; Platvoet et al. 2009), which would make it vulnerable to many different predators. Gammarids are an important prey for many fishes (Mazzi & Bakker 2003; Perrot-Minnot et al. 2007) and there are reports that native brown trout and perch can feed on killer shrimp in Britain (Aldridge 2015; Madgwick & Aldridge 2011). However, knowledge on the predators of killer shrimp is mostly anecdotal and there is little information on anti-predatory behaviour of this species in newly colonized areas, which is an important aspect to consider for predicting its future spread and impact.

Hiding, aggregation and crypsis are three of the most common anti-predatory strategies in aquatic species (Keenleyside 1979), which in the case of benthic gammarids are intimately related to the nature of the substrate (Holomuzki & Hoyle 1990). Hiding behaviour is particularly strong in gammarids (Goedmakers 1981; Jazdzewski et al. 2004), and availability of suitable substrate to hide can be a key determinant of establishment success in invasive gammarids (Devin et al. 2003), as different species may compete for shelter. For example, De Gelder et al. (2016) reported that the killer shrimp’s strong tendency to hide during daytime can displace the European native gammarid *Gammarus roeselii* from their shelters, which might put them at a higher risk of predation. Another common anti-predatory strategy is aggregation behaviour, as being part of a group can confuse predators (Krakauer 1995; Krause & Ruxton 2002) and reduce the per-capita probability of being preyed (Codella & Raffa 1995). Aggregation behaviour, however, also has costs as it is influenced by competition for food and mating partners, and poses a greater risk of being parasitized, which may put the group at a disadvantage (Krause & Ruxton 2002).

Thus, while killer shrimp invading Europe could be benefitting from a boom phase caused by predator release, the oddity effect and prey naïvety of novel predators might make them more vulnerable to native predators in the invaded waters. To shed light on this issue, we tested two anti-predatory behaviours (hiding and aggregation) in killer shrimp exposed to either dechlorinated water (control) or water conditioned with kairomones (i.e. semio-chemicals emitted by predators that allow eavesdropping by prey without benefitting the predator - Roberts & Garcia de Leaniz (2011) from a carnivorous fish predator, the three spined stickleback (*Gasterosteus aculeatus*). We wanted to test if killer shrimp from a recently colonized stream in Britain were able to recognise a common native fish as a predator or, on the contrary, displayed prey naïvety that might make it more difficult to mount and efficient anti-predatory response and, hence, make it more difficult to become established in neighbouring waters.

## Materials and Methods

### Collection and origin of samples

Killer shrimp (average size = 16.8 ± 0.9mm) were collected by live trapping in the Upper Mother Ditch (Margam, Wales, 51°33’19.5”N 3°44’46.6”W) in May 2017, and three-spined stickleback (weight range 0.9-2.0g) were hand-netted from an ornamental pond in Swansea (Singleton Park, Wales 51°36’26.2”N 3°58’52.4”W) in July 2017. We maintained the two species in separate 100L recirculation aquaculture systems at CSAR facilities (Swansea University) to avoid mixing their scents. Both species were fed frozen bloodworms, the sticklebacks every day and the killer shrimp three times per week. Water temperature was maintained at 15-16.5 °C with a weekly replacement of 20% volume.

### Experimental design

We set up two experiments to examine the killer shrimp’s anti-predatory behaviour in relation to the presence of stickleback’s kairomones (a fish predator that feeds on gammarids). In the first experiment, we compared the hiding behaviour of individual killer shrimp tested in water conditioned with stickleback kairomones compared to blank water. In the second experiment, we examined the attraction of single killer shrimp to the scent of conspecifics in an open-test arena scented with stickleback kairomones or with blank water. We chose the three-spined stickleback as a test predator because it is a common predator of gammarids (Macneil et al. 1999; Mazzi & Bakker 2003) and was present at the study site (Upper Mother Ditch) where killer shrimp were first detected in 2011, having been detected in a nearby reservoir one year earlier, in November 2010 (Madgwick & Aldridge 2011).

### Water conditioning

To obtain the kairomones used to simulate the presence of a predator, we housed 20 stickleback (biomass = 2.9 g/L) in a 10L tank of dechlorinated water for 24 hours. The conditioned water was prepared freshly the day before the experiments.

### Experiment 1. Hiding behaviour under the threat of predation

To quantify hiding behaviour under the threat of predation we used a 2L plastic tank (L:20cm, W:10cm, H:10cm) fitted with artificial grass patches (3cm^2^) glued to the bottom in a staggered fashion (Figure 1a), and a release cylinder (3.5cm diameter) located in the centre of the tank. At the beginning of the experiment 250ml of either dechlorinated water (control test) or fish-conditioned water (treatment) was added to the tank. One killer shrimp was placed inside the release cylinder and left to acclimatise for 5 min. The cylinder was then slowly lifted and the behaviour of the killer shrimp (time spent swimming or hiding in the artificial grass patches) was recorded for 10 minutes with a GoPro Hero camera mounted above the test tank (Figure 2A). In total, 20 individuals were tested with fish conditioned water and 20 individuals with blank water, the order of which was determined at random.

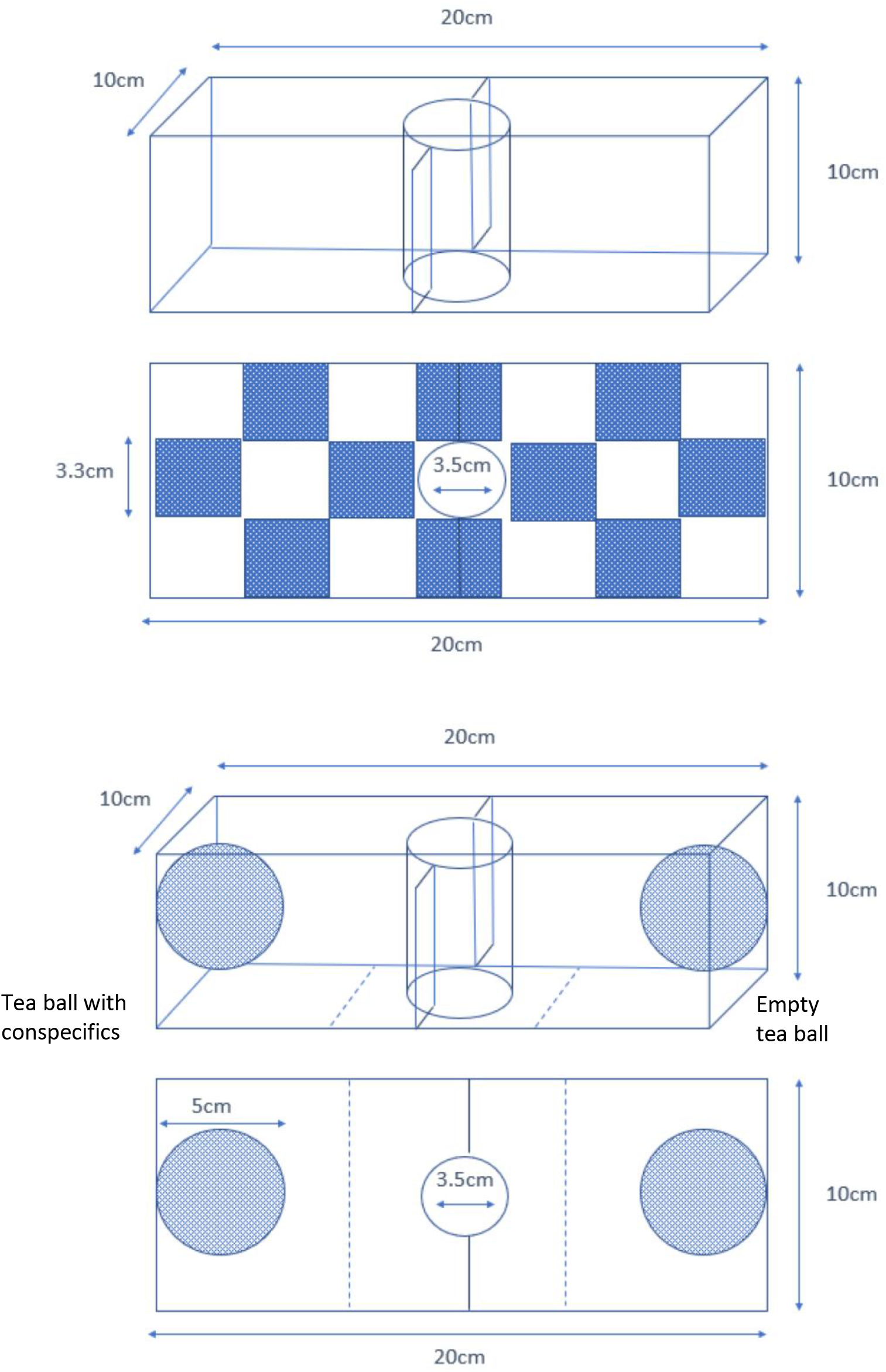

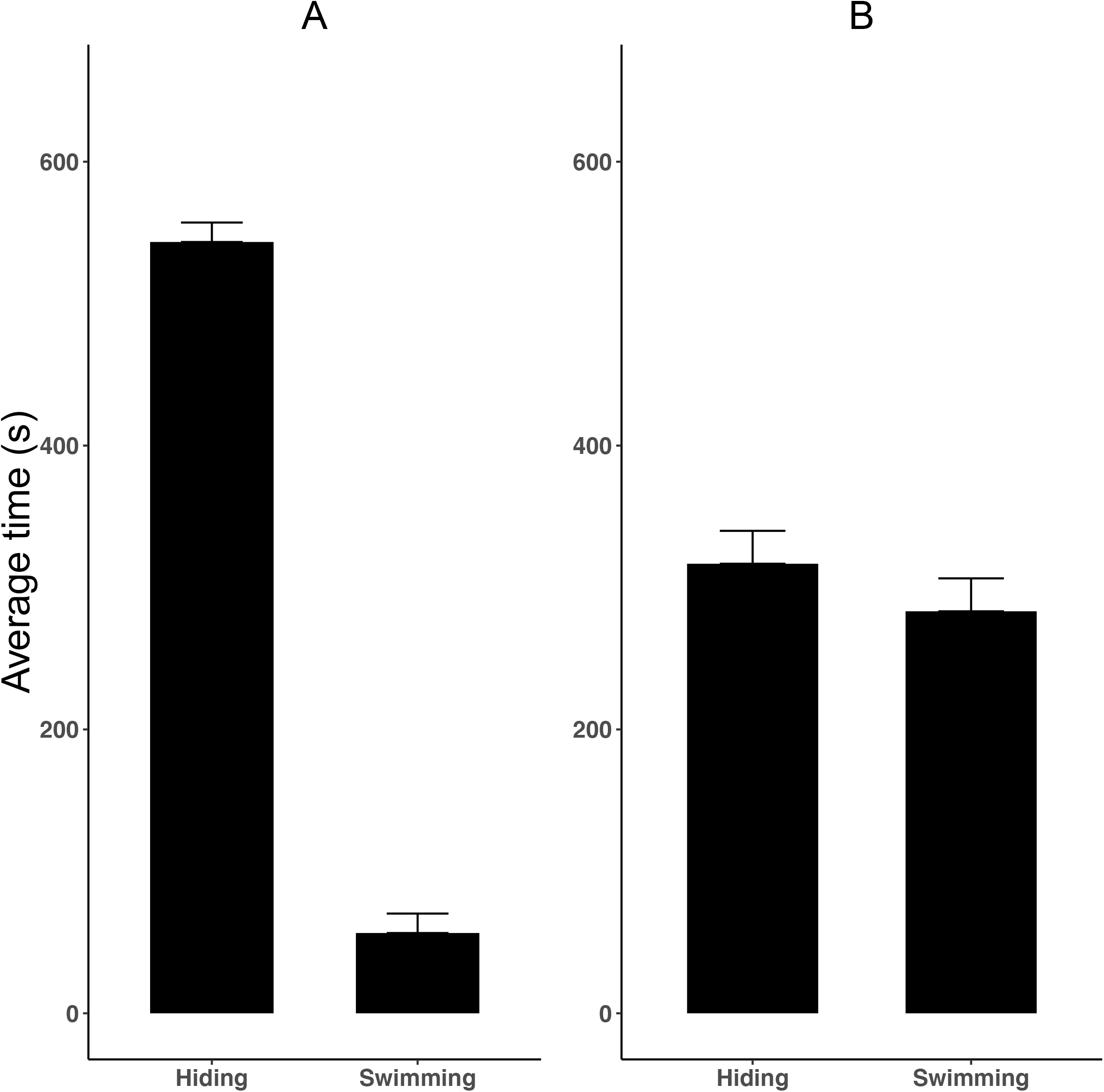

### Experiment 2. Conspecific attraction

To test if killer shrimp were more attracted to conspecifics under the threat of predation, we used a tank of the same size and volume (2 L) as the one used in experiment 1, but in this case the bottom was left bare and did not have artificial grass patches. At the two extremes of the tank we attached two tea balls (diameter 5cm) and drew two lines in the tank to notionally divide it into three equal sectors, two choice zones associated with the tea balls, and a middle section that served as a neutral (no choice) zone (Figure 1b). Ten killer shrimp were introduced in one of the two tea balls chosen at random, while the other one was left empty. As for experiment 1, 250ml of fish conditioned water or dechlorinated water were added to the tank and a single individual was introduced in the acclimatization cylinder, where it was left to acclimatize for 5min The cylinder was then removed, and the activity of the killer shrimp was recorded for 10 minutes with an overhead GoPro camera, as above. The time spent in each of the three tank zones was used to describe its behaviour: the time spent in the side containing the tea ball with conspecifics was interpreted as measure of attraction for group protection, the time spent in the central part was interpreted as neutral behaviour, and the time spent in the side with the empty tea ball was interpreted as avoidance of conspecifics. After each trial, the position of the two tea balls was alternated to control for possible external disturbances. In total, 40 killer shrimp were tested, 20 with dechlorinated water and 20 with fish scented water. The killer shrimp inside the tea ball were replaced between sessions to reduce aggressive behaviour due to confinement.

### Statistical analyses

We used R 3.3 (Team 2017), for analysis. For both experiments, we used a paired t-test to examine if (1) killer shrimp spent more time hiding than swimming when they were exposed to fish kairomones than when they were exposed to blank water (Experiment 1), and if (2) attraction to conspecifics was stronger when the killer shrimp were exposed to kairomones from a fish predator than when they were exposed to dechlorinated water (Experiment 2).

### Ethics Statement

Experiments were carried out in accordance with Swansea University’s Ethical guidelines and were approved by the Ethics Committee (070917/24, Reference Number: STU_BIOL_30638_060617140454_1). At the end of the experiments all sticklebacks were released alive at the site of capture. The killer shrimp, due to the risk they may pose for native communities, were disposed through incineration.

## Results

### Experiment 1. Hiding behaviour

Killer shrimp spent significantly more time hiding when the water was conditioned with kairomones from a predatory fish (mean time ± 95CI = 543.45 ± 13.7 s) than when they were tested against blank water (mean time ± 95CI = 386.75 ± 18.5 s; behaviour × treatment interaction *F*_1,76_ = 544.02, *P* < 0.001; Figure 2). Controls spent 50% of their time hiding and 50% swimming (*t*_19_ = 1.416, *P* = 0.173), whereas when they were exposed to fish kairomones they spent 91% of their time hiding and only 9% swimming (*t*_19_ = 34.789, *P* < 0.001).

### Experiment 2. Attraction to conspecifics

Killer shrimp spent much more time in the side of the tank scented with conspecifics (mean time 477.5 ± 20.5 s) than in the opposite side (mean time 39.1 ± 12.2 s), but such preference was not affected by the presence of fish kairomones (*t*_19_ = 0.245, *P* = 0.808; Figure 3).

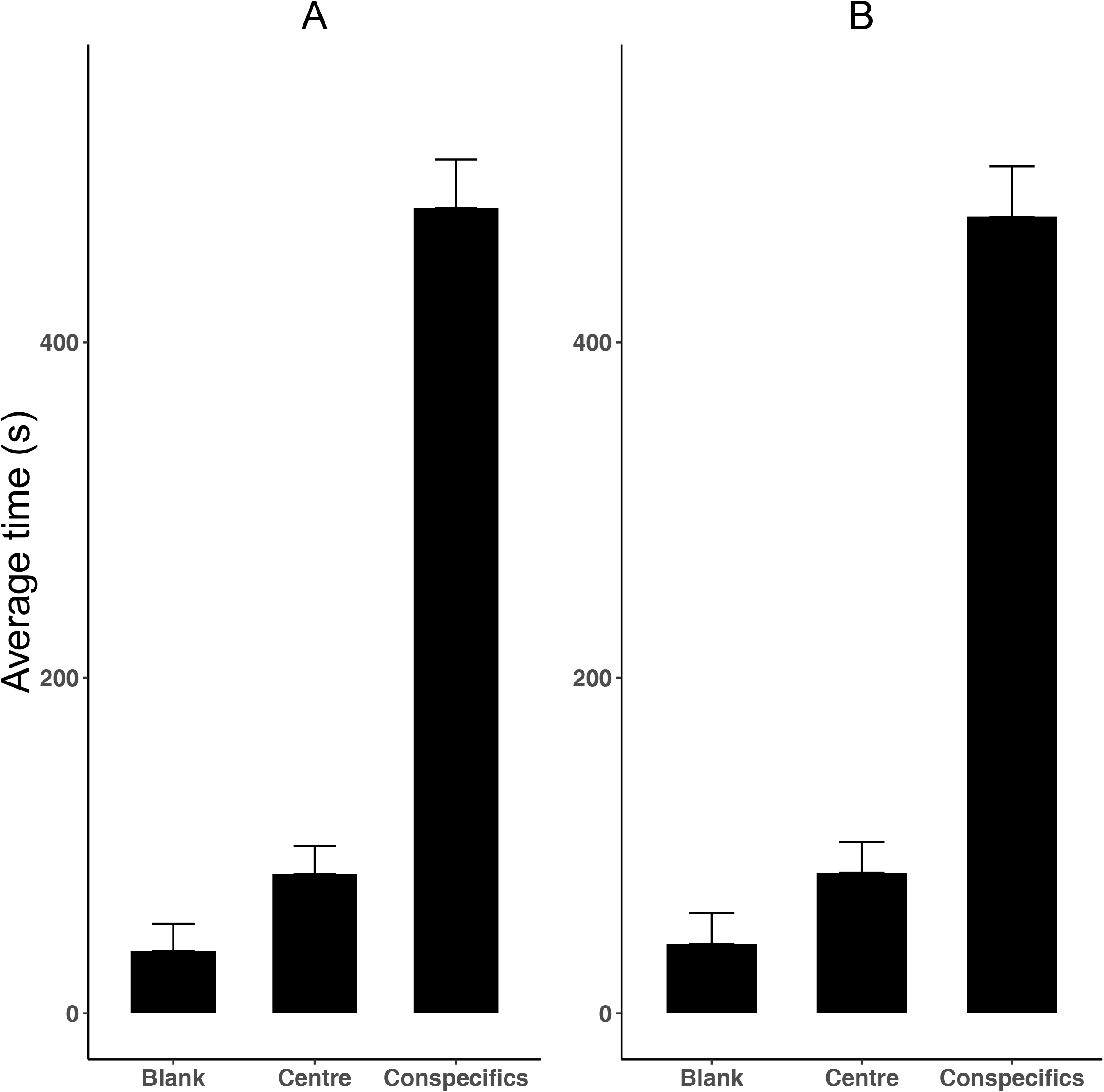

## Discussion

Our study shows that within 6 years (approximately 20 generations) of their introduction into a novel area in Britain killer shrimp display a strong tendency to hide when they are exposed to the chemical scent of a native predatory fish (the three spined stickleback), but not when they are exposed to dechlorinated water. Given that no evidence of predator avoidance was detected on the same population in relation to the scent of non-predatory Nile tilapia (Rolla et al. 2019), this strongly suggests that chemical recognition of stickleback kairomones constitutes an evolved, adaptive trait.

Much of our knowledge on the invasive killer shrimp refers to its role as a predator, there is little information regarding its role as a fish prey. This is unfortunate because predatory fish can exert strong selective pressures on gammarids (Åbjörnsson et al. 2004; Ahlgren et al. 2011; Kinzler & Maier 2006; Kotta et al. 2010; Wudkevich et al. 1997) and could play a major role in determining the killer shrimp’s invasion success. The killer shrimp has been found in the diet of 17 fish species found in the introduced range (9 exotic and 8 native, Table 1), but predator recognition has only been reported for the European bullhead *Cottus gobio* (Sornom et al. 2012), the racer goby *Babka gymnotrachelus* (Jermacz et al. 2017), and the spiny-cheek crayfish *Orconectes limosus* (Hesselschwerdt et al. 2009), therefore little is known about its antipredator behaviour. Amphipods can change their behaviour and habitat preferences when they detect chemical cues from potential predators (Baumgärtner et al. 2003; Thiel 2010), but also from injured conspecifics (Wisenden et al. 2001; Wudkevich et al. 1997), similar to what has been observed among teleost fishes (Roberts & Garcia de Leaniz 2011). Kairomone detection by gammarids has been reported previously as an anti-predatory strategy (Wudkevich et al. 1997), but seldom in the context of invasion biology (Hesselschwerdt et al. 2009).

Predation by native species could reduce, or at least slow down, invasions by non-native species (Zuharah & Lester 2010; Zuharah & Lester 2011) because they may not be able to recognise native predators (Sih et al. 2010), but also because their different appearance could make them easier to detect, or make them more attractive, to native predators (the ‘oddity prey effect’, (Penry□Williams et al. 2018). For example, killer shrimp are typically larger than native freshwater gammarids (Devin et al. 2003), and this might make it easier for visual predators to detect them. However, native predators may also be reluctant to feed on novel prey due to neophobia (Champneys et al. 2018), and this could result on lower predation pressure on invasive species (Crawley 1987; MacNeil et al. 2000; Trowbridge 1995; Wells & Henderson 1993). Killer shrimp could also benefit from a ‘shadow of safety’ effect if their relative low abundance during the earlier stages of invasion deflects predation pressure to the more abundant native prey (Trillo et al. 2016). Killer shrimp can rapidly become the dominant species in invaded benthonic communities (Dick & Platvoet 2000) and can become the most abundant food resource for fish feeding on macroinvertebrates. For example, field studies have indicated that killer shrimp can replace native *Gammarus roeseli* in the diet of zoo-benthivorous fish (Eckmann et al. 2008), but other studies have suggested the opposite, and reported that native fish prefer to feed on native gammarids (Kinzler & Maier 2006).

Clearly, the role of predation on invasion dynamics is difficult to predict, but knowledge of the time since introduction, and of prey-predator interactions appear important in determining establishment success. This is particularly complicated in the case of the killer shrimp in Great Britain because although its arrival is very recent, it may have already learned to chemically recognise a range of novel predators during its long invasion of Europe. Killer shrimp in the British Isles are genetically similar to those in continental Europe (Rewicz et al. 2015), where the invasion started in 1992 after the opening of the Main-Danube canal (Dick & Platvoet 2000), suggesting that they are stepping stones direct descendants from the first invaders. Stepping stones strategies drive the long distance dispersal of many species (Saura et al. 2014), and it is possible that repeated residencies in different habitats may have enabled the killer shrimp to learn to recognise different predators. Given that the three-spined stickleback is also widespread in continental Europe, our study cannot rule out that the observed predator recognition was acquired in Britain, or represents an older behavioural legacy from previous invasions.

Two common anti-predatory strategies in amphipods are to reduce mobility and become more aggregated under the risk of predation (Åbjörnsson et al. 2000; Williams & Moore 1985; Williams et al. 2016). Results from Experiment 1 in our study indicate that killer shrimp spend more time hiding and less time swimming when they were exposed to predator kairomones, as seen in other gammarids. These findings are also in agreement with those of Sornom et al. (2012) who observed a decrease in mobility and an increase in hiding time in killer shrimp exposed to the scent of another fish predator, the European bullhead (*Cottus gobio*). However, our results on aggregation behaviour (Experiment 2) are more equivocal. Unlike *Gammarus pulex*, which become increasingly aggregated when exposed to stickleback kairomones (Kullmann et al. 2008), killer shrimp in our study showed the same strong preference to remain in the vicinity of conspecifics even when there was no immediate threat of predation. Exposure to bullhead kairomones also failed to elicit an increase in killer shrimp aggregation (Sornom et al. 2012), but in this case aggregation was low. Jermacz et al. (2017) have shown that killer shrimp prefer to hide in response to predator cues, rather than aggregate, when refuges are present, and that they aggregate when there are no shelters and staying in a group is the only antipredator strategy possible. It is possible that aggregation behaviour in the killer shrimp depends on the availability of shelters, but also on the risk of intra-guild predation. Compared to native gammarids, killer shrimp display higher sociability and lower incidence of cannibalism (Kinzler et al. 2009; Truhlar & Aldridge 2015), which may explain their strong tendency to aggregate. Aggregation behaviour can provide not only protection from fish predators (Åbjörnsson et al. 2004), but could also facilitate dispersal, as living in a group would increase the number of founders, and propagule pressure has been found to be an important factor determining invasion success (Consuegra et al. 2011; Ricciardi et al. 2010)

## Conclusions

In conclusion, prey-predator dynamics are an important, but largely neglected, determinants of invasion success and our study indicates that knowledge of anti-predatory strategies might be important for predicting dispersal pathways and risk of establishment in the killer shrimp, and likely also on other aquatic invaders. Killer shrimp are dispersing at an alarmingly fast rate in Europe (DAISIE 2009; Gallardo et al. 2016), and prevention and control measures might benefit from information on prey and predators present in communities at risk. In this sense, behavioural profiling of anti-predatory strategies, using perhaps some of the simple assays shown in our study, could be incorporated into risk assessments. Knowledge of how invasive species might respond to resident predators can inform the development of more efficient management actions, as these seldom consider biotic resistance (Robinson et al. 2019; Robinson et al. 2018). Given its strong aggregation behaviour, we also suggest that even when complete eradication is not possible, control measures that aim to reduce the density of killer shrimp might be beneficial, as a lower relative abundance and a smaller group size can make them more vulnerable to fish predators, potentially reducing their impact on native communities.

## Supporting information

Fish predators of killer shrimp

## Acknowledgments

We are grateful to Emma Keenan and Graham Rutt (Natural Resources Wales) for help with the sampling. Funding was provided by the EC Horizon 2020 Aquainvad-ED project (Marie Sklodowska-Curie ITN-2014-ETN-642197) to SC.

## Author Contributions & Competing Interests

MR and CGL designed the study. SC and CGL secured the funding. MR collected the data and carried out the analyses with advice from CGL. MR and CGL wrote the MS with contributions from SC. The authors declare no competing interests.

## References

Åbjörnsson K, Dahl J, Nyström P, and Brönmark C. 2000. Influence of predator and dietary chemical cues on the behaviour and shredding efficiency of Gammarus pulex. Aquatic Ecology 34:379–387.

Åbjörnsson K, Hansson L-A, and Brönmark C. 2004. Responses of prey from habitats with different predator regimes: local adaptation and heritability. Ecology 85:1859–1866.

Ahlgren J, Åbjörnsson K, and Brönmark C. 2011. The influence of predator regime on the behaviour and mortality of a freshwater amphipod, Gammarus pulex. Hydrobiologia 671:39–49.

Aldridge DC. 2015. Killer shrimp, Dikerogammarus villosus. Sand Hutton, York, UK: GB Non-native Species Secretariat

Almany GR, Peacock LF, Syms C, McCormick MI, and Jones GP. 2007. Predators target rare prey in coral reef fish assemblages. Oecologia 152:751–761.

Arundell K, Dunn A, Alexander J, Shearman R, Archer N, and Ironside JE. 2015. Enemy release and genetic founder effects in invasive killer shrimp populations of Great Britain. Biological Invasions 17:1439–1451.

Bacela-Spychalska K, and van der Velde G. 2013. There is more than one ‘killer shrimp’: trophic positions and predatory abilities of invasive amphipods of Ponto-Caspian origin. Freshwater Biology 58:730–741.

Baumgärtner D, Koch U, and Rothhaupt K-O. 2003. Alteration of kairomone-induced antipredator response of the freshwater amphipod Gammarus roeseli by sediment type. Journal of chemical ecology 29:1391–1401.

Borcherding J, Dolina M, Heermann L, Knutzen P, Krüger S, Matern S, van Treeck R, and Gertzen S. 2013. Feeding and niche differentiation in three invasive gobies in the Lower Rhine, Germany. Limnologica-Ecology and Management of Inland Waters 43:49–58.

Casellato S, Visentin A, and La Piana G. 2007. The predatory impact of Dikerogammarus villosus on fish. Biological invaders in inland waters: profiles, distribution, and threats: Springer, 495–506.

Champneys T, Castaldo G, Consuegra S, and Garcia de Leaniz C. 2018. Density-dependent changes in neophobia and stress-coping styles in the world’s oldest farmed fish. Royal Society Open Science 5:181473. doi:10.1098/rsos.181473

Codella SG, and Raffa KF. 1995. Contributions of female oviposition patterns and larval behavior to group defense in conifer sawflies (Hymenoptera: Diprionidae). Oecologia 103:24–33.

Colautti RI, Ricciardi A, Grigorovich IA, and MacIsaac HJ. 2004. Is invasion success explained by the enemy release hypothesis? Ecology Letters 7:721–733.

Consuegra S, Phillips N, Gajardo G, and Garcia de Leaniz C. 2011. Winning the invasion roulette: Escapes from fish farms increase admixture and facilitate establishment of non-native rainbow trout. Evolutionary Applications 4:660–671.

Crawley MJ. 1987. What makes a community invasible? Colonization, succession and stability:429–453.

DAISIE. 2009. Handbook of alien species in Europe. Dordrecht: Springer.

De Gelder S, Van der Velde G, Platvoet D, Leung N, Dorenbosch M, Hendriks H, and Leuven R. 2016. Competition for shelter sites: Testing a possible mechanism for gammarid species displacements. Basic and applied ecology 17:455–462.

Devin S, and Beisel J-N. 2008. Geographic patterns in freshwater gammarid invasions: an analysis at the pan-European scale. Aquatic Sciences 70:100–106.

Devin S, Piscart C, Beisel J-N, and Moreteau J. 2003. Ecological traits of the amphipod invader Dikerogammarus villosus on a mesohabitat scale. Archiv für Hydrobiologie 158:43–56.

Dick JT, and Platvoet D. 2000. Invading predatory crustacean Dikerogammarus villosus eliminates both native and exotic species. Proceedings of the Royal Society of London Series B: Biological Sciences 267:977–983.

Djikanovic V, Gacic Z, and Cakic P. 2010. Endohelminth fauna of barbel Barbus barbus (L. 1758) in the Serbian section of the Danube River, with dominance of acanthocephalan *Pomphorhynchus laeavis*. Bulletin of the European Association of Fish Pathologists 30:229–236.

Eckmann R, Mörtl M, Baumgärtner D, Berron C, Fischer P, Schleuter D, and Weber A. 2008. Consumption of amphipods by littoral fish after the replacement of native Gammarus roeseli by invasive Dikerogammarus villosus in Lake Constance. Aquatic Invasions 3:187–191.

Gallardo B, Zieritz A, Adriaens T, Bellard C, Boets P, Britton JR, Newman JR, van Valkenburg JLCH, and Aldridge DC. 2016. Trans-national horizon scanning for invasive non-native species: a case study in western Europe. Biological Invasions 18:17–30. 10.1007/s10530-015-0986-0

Giller PS. 2000. Dominant role of exotic invertebrates, mainly Crustacea. The Biodiversity Crisis and Crustacea-Proceedings of the Fourth International Crustacean Congress: CRC Press. p 35.

Goedmakers A. 1981. Population dynamics of three gammarid species. Bijdragen tot de Dierkunde 51:181–190.

Grabner DS, Weigand AM, Leese F, Winking C, Hering D, Tollrian R, and Sures B. 2015. Invaders, natives and their enemies: distribution patterns of amphipods and their microsporidian parasites in the Ruhr Metropolis, Germany. Parasites & vectors 8:419.

Haubrock PJ, Balzani P, Johovic I, Inghilesi AF, Nocita A, and Tricarico E. 2018. The diet of the alien channel catfish Ictalurus punctatus in the River Arno (Central Italy). Aquatic Invasions 13.

Hesselschwerdt J, Tscharner S, Necker J, and Wantzen KM. 2009. A local gammarid uses kairomones to avoid predation by the invasive crustaceans Dikerogammarus villosus and Orconectes limosus. Biological Invasions 11:2133.

Holomuzki JR, and Hoyle JD. 1990. Effect of predatory fish presence on habitat use and diel movement of the stream amphipod, *Gammarus minus*. Freshwater Biology 24:509–517.

Jazdzewski K, Konopacka A, and Grabowski M. 2004. Recent drastic changes in the gammarid fauna (Crustacea, Amphipoda) of the Vistula River deltaic system in Poland caused by alien invaders. Diversity and Distributions 10:81–87.

Jermacz Ł, Andrzejczak J, Arczyńska E, Zielska J, and Kobak J. 2017. An enemy of your enemy is your friend: Impact of predators on aggregation behavior of gammarids. Ethology 123:627–639.

Jurajda P, Všetičková L, Polačik M, and Vassilev M. 2013. Can round goby (*Neogobius melanostomus*) caught by rod and line be used for diet analysis? Journal of Great Lakes Research 39:182–185.

Keenleyside MH. 1979. Anti-predator behaviour. Diversity and Adaptation in Fish Behaviour: Springer, 44–62.

Kinzler W, Kley A, Mayer G, Waloszek D, and Maier G. 2009. Mutual predation between and cannibalism within several freshwater gammarids: Dikerogammarus villosus versus one native and three invasives. Aquatic Ecology 43:457.

Kinzler W, and Maier G. 2006. Selective predation by fish: a further reason for the decline of native gammarids in the presence of invasives? Journal of Limnology 65:27–34.

Kotta J, Orav-Kotta H, and Herkül K. 2010. Separate and combined effects of habitat-specific fish predation on the survival of invasive and native gammarids. Journal of Sea Research 64:369–372.

Krakauer DC. 1995. Groups confuse predators by exploiting perceptual bottlenecks: a connectionist model of the confusion effect. Behavioral Ecology and Sociobiology 36:421–429.

Krause J, and Ruxton GD. 2002. Living in groups: Oxford University Press.

Kullmann H, Thünken T, Baldauf SA, Bakker TC, and Frommen JG. 2008. Fish odour triggers conspecific attraction behaviour in an aquatic invertebrate. Biol Lett 4:458–460.

Maazouzi C, Médoc V, Pihan J-C, and Masson G. 2011. Size-related dietary changes observed in young-of-the-year pumpkinseed (*Lepomis gibbosus*): stomach contents and fatty acid analyses. Aquatic Ecology 45:23–33.

Macneil C, Boets P, Lock K, and Goethals PLM. 2013. Potential effects of the invasive ‘killer shrimp’ (Dikerogammarus villosus) on macroinvertebrate assemblages and biomonitoring indices. Freshwater Biology 58:171–182. 10.1111/fwb.12048

MacNeil C, Boets P, and Platvoet D. 2012. ‘Killer shrimps’, dangerous experiments and misguided introductions: how freshwater shrimp (Crustacea: Amphipoda) invasions threaten biological water quality monitoring in the British Isles. Freshwater Reviews 5:21–35. 10.1608/frj-5.1.457

Macneil C, Dick JT, and Elwood RW. 1999. The dynamics of predation on Gammarus spp.(Crustacea: Amphipoda). Biological Reviews 74:375–395.

MacNeil C, Elwood RW, and Dick JT. 2000. Factors influencing the importance of Gammarus spp.(Crustacea: Amphipoda) in riverine salmonid diets. Archiv für Hydrobiologie:87–107.

Madgwick G, and Aldridge DC. 2011. Killer shrimps in Britain: hype or horror. British Wildllife 22:408–412.

Mayer G, Maier G, Maas A, and Waloszek D. 2008. Mouthparts of the ponto-caspian invader Dikerogammarus villosus (Amphipoda: Pontogammaridae). Journal of Crustacean Biology 28:1–15.

Mazzi D, and Bakker T. 2003. A predator’s dilemma: prey choice and parasite susceptibility in three-spined sticklebacks. Parasitology 126:339–347.

Penry□Williams IL, Ioannou CC, and Taylor MI. 2018. The oddity effect drives prey choice but not necessarily attack time. Ethology 124:496–503.

Perrot-Minnot M-J, Kaldonski N, and Cézilly F. 2007. Increased susceptibility to predation and altered anti-predator behaviour in an acanthocephalan-infected amphipod. International Journal for Parasitology 37:645–651.

Platvoet D, Van Der Velde G, Dick JT, and Li S. 2009. Flexible omnivory in Dikerogammarus villosus (Sowinsky, 1894)(Amphipoda)-amphipod pilot species project (AMPIS) Report 5. Crustaceana:703–720.

Pöckl M. 2009. Success of the invasive Ponto-Caspian amphipod Dikerogammarus villosus by life history traits and reproductive capacity. Biological Invasions 11:2021–2041.

Rewicz T, Wattier R, Grabowski M, Rigaud T, and Bącela-Spychalska K. 2015. Out of the Black Sea: phylogeography of the invasive killer shrimp Dikerogammarus villosus across Europe. PloS one 10:e0118121.

Ricciardi A, Jones LA, Kestrup ÅSM, and Ward JM. 2010. Expanding the propagule pressure concept to understand the impact of biological invasions. In: Richardson DM, ed. Fifty Years of Invasion Ecology: Wiley-Blackwell, 225–238.

Roberts LJ, and Garcia de Leaniz C. 2011. Something smells fishy: predator-naïve salmon use diet cues, not kairomones, to recognize a sympatric mammalian predator. Animal Behaviour 82:619–625. 10.1016/j.anbehav.2011.06.019

Robinson CV, Garcia de Leaniz C, and Consuegra S. 2019. Effect of artificial barriers on the distribution of the invasive signal crayfish and Chinese mitten crab Scientific Reports.

Robinson CV, Garcia de Leaniz C, James J, Cable J, Orozco-terWengel P, and Consuegra S. 2018. Genetic diversity and parasite facilitated establishment of the invasive signal crayfish (*Pacifastacus leniusculus*) in Great Britain. Ecology and Evolution 8:9181–9191. 10.1002/ece3.4235

Rolla M, Consuegra S, Carrington E, Hall D, and Garcia de Leaniz C. 2019. Experimental evidence of invasion facilitation in the zebra mussel-killer shrimp system. bioRxiv:626432. 10.1101/626432

Saura S, Bodin Ö, and Fortin MJ. 2014. Stepping stones are crucial for species’ long-distance dispersal and range expansion through habitat networks. Journal of Applied Ecology 51:171–182.

Semenov DY. 2009. Data on morphology and biology of Caspian big-headed goby *Neogobius gorlap* (Perciformes, Gobiidae) from the Kuibyshev Reservoir. Journal of Ichthyology 49:834–837.

Sih A, Bolnick DI, Luttbeg B, Orrock JL, Peacor SD, Pintor LM, Preisser E, Rehage JS, and Vonesh JR. 2010. Predator–prey naïveté, antipredator behavior, and the ecology of predator invasions. Oikos 119:610–621.

Sornom P, Gismondi E, Vellinger C, Devin S, Férard J-F, and Beisel J-N. 2012. Effects of sublethal cadmium exposure on antipredator behavioural and antitoxic responses in the invasive amphipod *Dikerogammarus villosus*. PloS one 7:e42435.

Strayer DL, D’Antonio CM, Essl F, Fowler MS, Geist J, Hilt S, Jaric I, Johnk K, Jones CG, Lambin X, Latzka AW, Pergl J, Pysek P, Robertson P, von Schmalensee M, Stefansson RA, Wright J, and Jeschke JM. 2017. Boom-bust dynamics in biological invasions: towards an improved application of the concept. Ecol Lett 20:1337–1350. 10.1111/ele.12822

Team RC. 2017. R: A language and environment for statistical computing Vienna, Austria: R Foundation for Statistical Computing.

Thiel M. 2010. Chemical communication in peracarid crustaceans. Chemical communication in crustaceans: Springer, 199–218.

Tricarico E, Mazza G, Orioli G, Rossano C, Scapini F, and Gherardi F. 2010. The killer shrimp, *Dikerogammarus villosus* (Sowinsky, 1894), is spreading in Italy. Aquatic Invasions 5:211–214. 10.3391/ai.2010.5.2.14

Trillo PA, Bernal XE, Caldwell MS, Halfwerk WH, Wessel MO, and Page RA. 2016. Collateral damage or a shadow of safety? The effects of signalling heterospecific neighbours on the risks of parasitism and predation. Proceedings of the Royal Society B: Biological Sciences 283:20160343.

Trowbridge CD. 1995. Establishment of the green alga Codium fragile ssp. tomentosoides on New Zealand rocky shores: current distribution and invertebrate grazers. Journal of Ecology:949–965.

Truhlar AM, and Aldridge DC. 2015. Differences in behavioural traits between two potentially invasive amphipods, Dikerogammarus villosus and Gammarus pulex. Biological Invasions 17:1569–1579.

Wells JD, and Henderson G. 1993. Fire ant predation on native and introduced subterranean termites in the laboratory: effect of high soldier number in *Coptotermes formosanus*. Ecological Entomology 18:270–274.

Williams DD, and Moore KA. 1985. The role of semiochemicals in benthic community relationships of the lotic amphipod *Gammarus pseudolimnaeus*: a laboratory analysis. Oikos:280–286.

Williams KL, Navins KC, and Lewis SE. 2016. Behavioral responses to predation risk in brooding female amphipods (Gammarus pseudolimnaeus). Journal of Freshwater Ecology 31:571–581.

Wisenden BD, Pohlman SG, and Watkin EE. 2001. Avoidance of conspecific injury-released chemical cues by free-ranging *Gammarus lacustris* (Crustacea: Amphipoda). Journal of chemical ecology 27:1249–1258.

Wudkevich K, Wisenden BD, Chivers DP, and Smith RJF. 1997. Reactions of *Gammarus lacustris* to chemical stimuli from natural predators and injured conspecifics. Journal of chemical ecology 23:1163–1173.

Zuharah WF, and Lester PJ. 2010. Can adults of the New Zealand mosquito *Culex pervigilans* (Bergorth) detect the presence of a key predator in larval habitats? Journal of Vector Ecology 35:100–105.

Zuharah WF, and Lester PJ. 2011. Are exotic invaders less susceptible to native predators? A test using native and exotic mosquito species in New Zealand. Population Ecology 53:307–317.

