## Supplementary material for "Predator recognition and anti-predatory behaviour in a recent aquatic invader, the killer shrimp (*Dikerogammarus villosus*)": Fish predators of killer shrimp

**Table 1.** Fish predators of killer shrimp.

| **Family** | **Species** | **Status** | **Recognition** | **Location** | **Reference** |
| --- | --- | --- | --- | --- | --- |
| Gobiidae | Caspian big headed goby  *Neogobius gorlap* | Exotic | NA | Kuibyshev Reservoir (Russia) | Semenov (2009) |
|  | Round goby  *Neogobius elanostomus* | Exotic | NA | Danube River (Bulgaria) | Jurajda et al. (2013) |
|  | Monkey goby  *Neogobius fluviatilis* | Exotic | NA | River Rhine (Germany) | Borcherding et al. (2013) |
|  | bighead goby  *Ponticola kessleri* | Exotic | NA | River Rhine (Germany) | Borcherding et al. (2013) |
|  | Racer goby  *Babka gymnotrachelus* | Exotic | Y | Laboratory (Poland) | Jermacz et al. (2017) |
| Percidae | Eurasian ruffe  *Gymnocephalus cernua* | Native | NA | River Rhine (Netherlands) | Giller (2000) |
|  | Zander  *Sander lucioperca* | Native | NA | River Rhine (Netherlands) | Giller (2000) |
|  | Eurasian perch  *Perca fluviatilis* | Native | NA | Constance Lake (Germany, Austria, Switzerland) | Eckmann et al. (2008) |
|  | Eurasian perch  *Perca fluviatilis* | Native | NA | Grafham Reservoir (UK) | Madgwick & Aldridge (2011) |
| Salmonidae | Brown trout  *Salmo trutta* | Native | NA | Grafham Reservoir (UK) | Madgwick & Aldridge (2011) |
|  | Rainbow trout  *Oncorhynchus mykiss* | Exotic | NA | Grafham Reservoir (UK) | Madgwick & Aldridge (2011) |
| Cottidae | European bullhead  *Cottu*s *gobio* | Native | Y | Laboratory (France) | Sornom et al. (2012) |
| Lotidae | Burbot  *Lota lota* | Native | NA | Constance Lake (Germany, Austria, Switzerland) | Eckmann et al. (2008) |
| Anguillidae | European eel  *Anguilla anguilla* | Native | NA | Constance Lake (Germany, Austria, Switzerland) | Eckmann et al. (2008) |
| Cyprinidae | Common barbel  *Barbus barbus* | Native | NA | Danube River (Serbia) | Djikanovic et al. (2010) |
| Centrarchidae | Pumpkinseed  *Lepomis gibbosus* | Exotic | NA | Mirgenbach Reservoir (France) | Maazouzi et al. (2011) |
| Ictaluridae | Channel catfish  *Ictalurus punctatus­* | Exotic | NA | River Arno (Italy) | Haubrock et al. (2018) |
| Siluridae | Wels catfish  *Silurus glanis* | Exotic | NA | River Arno (Italy) | Haubrock et al. (2018) |
